# Stochastic to Deterministic: A straightforward approach to create serially perfusable multiscale capillary beds

**DOI:** 10.1101/2024.05.03.592474

**Authors:** Michael J. Donzanti, Omkar Mhatre, Brea Chernokal, Diana C. Renteria, Jason P. Gleghorn

## Abstract

Generation of *in vitro* tissue models with serially perfused hierarchical vasculature would allow greater control of fluid perfusion throughout the network and enable direct mechanistic investigation of vasculogenesis, angiogenesis, and vascular remodeling. In this work, we have developed a method to produce a closed, serially perfused, multiscale vessel network embedded within an acellular hydrogel. We confirmed that the acellular and cellular gel-gel interface was functionally annealed without preventing or biasing cell migration and endothelial self-assembly. Multiscale connectivity of the vessel network was validated via high-resolution microscopy techniques to confirm anastomosis between self-assembled and patterned vessels. Lastly, using fluorescently labeled microspheres, the multiscale network was serially perfused to confirm patency and barrier function. Directional flow from inlet to outlet mandated flow through the capillary bed. This method for producing closed, multiscale vascular networks was developed with the intention of straightforward fabrication and engineering techniques so as to be a low barrier to entry for researchers who wish to investigate mechanistic questions in vascular biology. This ease of use offers a facile extension of these methods for incorporation into organoid culture, organ-on-a-chip (OOC) models, and bioprinted tissues.

## Introduction

Creating vascularized tissue has been a longstanding goal of bioengineering to enable large-scale tissue engineering and vascular biology studies (1–3). As such numerous approaches have been developed to generate *in vitro* vascular tissue models. Methods to create large vessels often rely on “top-down” deterministic approaches, including micro-molding via casts and needles, direct 3D printing of cellular biomaterials to create vessel channels, or 3D printing of multi-ink systems including sacrificial biomaterials to form a vessel lumen (3–18). However, due to fabrication resolution limitations, these methods cannot be used to generate small vessels, so microvasculature is often created via endothelial self-assembly (19–27) which generates a vascular network called a plexus. Perfusable plexuses are most often fabricated as small regions within microfluidic devices; however, recent work has demonstrated the capability to create large perfusable self-assembled plexuses with tunable vessel sizes within arbitrarily shaped collagen gels (28). Notably, a common issue with endothelial self-assembly outside of a microfluidic device is that the network is not closed but instead has open, “leaky” vessels at the boundaries of the gel.

This poses a problem not only for controllable perfusion but also for incorporation into a hierarchical vascular network. These two features are necessary prerequisites for any viable *in vitro* vascular tissue model aimed at investigating vascular development and remodeling. Simple methods to generate closed mixed vessel sizes in a hierarchically perfused vascular system have not yet been realized due to these fabrication limitations.

The combination of multiple length scales has been historically difficult to achieve without the need for specialized device housings for successful perfusion (3, 29, 30). The standard approach used to perfuse an *in vitro* microvascular network incorporates large media channels that run parallel to a self-assembled endothelial network embedded within a central hydrogel (20, 31–39). Indeed, these designs are sufficient engineering strategies for the perfusion of self-assembled networks. Flow into the network can be directed via tuning the large channel flow rates and these channels serve as boundaries to the “leaky” vascular network at the hydrogel edge. However, the design of these systems results in parallel fluid flow throughout the entire self-assembled network. Whereas this configuration results in a functional, perfusable, self-assembled microvascular network, these geometries do not recapitulate vasculature *in vivo*, which is organized into a hierarchical network that provides serial perfusion of distal microvasculature.

To investigate vascular development, refinement, and remodeling, simple and reliable methods need to be created to generate multiscale vascular networks that are hierarchically perfused. It is well established that tissue developmental processes are mechanically regulated by a number of cues, including fluid flow which is tightly linked to vessel network geometry (40–43). Fluid shear stress is a critical regulator of endothelial function and organization. Indeed, fluid shear stress has been shown to play a significant role in angiogenesis in multiple vascular models, including within patterned vessels (44, 45); however, the role of fluid forces in self-assembled networks and their remodeling into organized vasculature is still an emerging area of study *in vitro* (46). In addition, the hierarchical nature of vasculature *in vivo* allows for the creation of growth factor gradients and convection of secreted factors from proximal vessels to distal microvascular beds. Generation of *in vitro* tissue models with serially perfused hierarchical vasculature would allow greater control of fluid perfusion throughout the network and enable direct mechanistic investigation of vasculogenesis, angiogenesis, and vascular remodeling.

As such, we have developed a method for the incorporation of multiscale vasculature into a standalone construct by embedding a cellular self-assembled network within an acellular collagen gel. Large deterministically patterned vessels are connected in series to the stochastic, self-assembled microvascular network, thereby requiring fluid flow through the capillary bed for complete perfusion. The configuration of this perfused capillary bed allows for integration with multiple fabrication methods to generate deterministically patterned or printed large vessel geometries with fixed inlets and outlets to enable straightforward connection to external perfusion pumps if desired. Serial flow through the hierarchical structure more faithfully recapitulates native *in vivo* vascular architecture, allowing for the investigation of fluid flows and mechanotransductive signals at the level of a vascular network as opposed to an individual vessel. Additionally, this system was specifically designed to be fabricated straightforwardly, without specialized fabrication equipment, to increase the application space of these engineered capillary beds into organoid culture, organ-on-a-chip (OOC) models, and bioprinted tissues. Ideally, these approaches can act as a building block for micro-macro integrated systems to generate a variety of hierarchically organized tissue systems (47). This strategy of building serial, hierarchically organized vascular networks is a powerful tool for vascular morphogenesis and mechanistic vascular biology studies that are not possible with current approaches.

## Materials and Methods

### Reagents

#### Rat tail collagen isolation

Methods of collagen isolation were similar to previous studies (24, 28). Tendons were removed from rat tails (Pel-Freez Biologicals, Rogers, AR) and soaked in 1X DPBS (Corning, Corning, NY). Tendons were transferred to acetone for 5 min, followed by 70% isopropanol for 5 min. Tendons were then transferred to 0.1% glacial acetic acid and dissolved for 48 hours at 4°C. After 48 hours, the solution was centrifuged at 28,000 x g for 1 hour. The supernatant was removed, frozen at -80°C, and lyophilized. The lyophilized collagen was dissolved in 0.1% glacial acetic acid at an 8 mg/mL stock concentration and stored at 4°C for further use.

#### Transduction and culture of HUVECs

Human umbilical vein endothelial cells (HUVECs, Lonza, Basel, CH) were cultured on a 0.1% gelatin-coated tissue culture dish at 37°C and 5% CO_2_ in Endothelial Cell Growth Medium - 2(EGM-2, Lonza, Basel, CH). HUVECs were transduced with mCherry lentiviral pseudovirus (pCDH-CMV-mCherry-T2A-Puro, Addgene, Watertown, MA) to generate fluorescently labeled cells at approximately 60% confluency for 24h. Transduced HUVECs were positively selected and maintained over culture with 0.1% puromycin (VWR International, Radnor, PA). All cells used in these studies were from passages six to nine.

#### Cellular collagen gel and culture medium

For HUVEC-laden gels, HUVECs were lifted with 0.05% trypsin-EDTA (Corning, Corning, NY), pelleted at 300 × g for 5 min, and resuspended in supplemented medium. Collagen gel stock was neutralized with NaOH, diluted to the working concentration of 3 mg/mL, as previously described (24), and the cell suspension was added to result in a final cell concentration of 1x10^6^ cells/mL of gel. Culture medium used for HUVEC cellular gels consisted of 2% vasculogenesis medium (EGM-2 medium supplemented with 50 ng/mL tetradecanoylphorbol acetate (TPA, Adipogen, San Diego, CA) and 50 μg/mL sodium ascorbate). For fibroblast-seeded gels, NIH-3T3 cells (ATCC, Gaithersburg, MD) were added to neutralized and diluted 3 mg/mL collagen solution, as described above, at a density of 1x10^6^ cells/mL of gel. Culture medium for used 3T3 cellular gels consisted of DMEM (Corning, Corning, NY) supplemented with 10% fetal bovine serum (ThermoFisher, Waltham, MA) and 1% penicillin/streptomycin (ThermoFisher, Waltham, MA). Gels without cells were made by neutralizing the collagen solution with NaOH then diluting the solution to the desired collagen concentration.

### Characterization of collagen-collagen gel interfaces

#### Creation of collagen-collagen gels

Collagen gels were fabricated using techniques similar to previous work (31, 32, 48–56). Square, 1 cm x 1 cm wells were cut from a polydimethylsiloxane (PDMS; Sylgard Dow Corning, Midland, MI)) sheet prior to bonding to a glass slide with oxygen plasma. A PDMS barrier was inserted on one side of the well to form two side-by-side compartments. The first collagen prepolymer was seeded into the open compartment and allowed to polymerize for a specified time (30 s, 1 min, 10 min, 30 min). The PDMS barrier was then quickly removed, and the remaining volume of the well was seeded with a second collagen prepolymer solution. Gel constructs were allowed to fully polymerize at 37°C for 20 min before DPBS was added to prevent drying. Devices were kept in PBS at room temperature until imaging. PDMS culture wells were used to fabricate acellular collagen-collagen constructs or cellular-acellular collagen-collagen constructs. For constructs with cells, either HUVECs or 3T3s were seeded into a collagen prepolymer as described above, and the same protocols were followed with the addition of cell culture medium for the corresponding cell type in lieu of PBS after gel formation. Cellular gel constructs were cultured at 37°C with 5% CO_2_ for 7 days with daily media changes.

#### Imaging and characterization of collagen-collagen gels

Acellular collagen-collagen devices were imaged via reflectance confocal microscopy (10X, LSM880, Zeiss) to visualize collagen density and orientation. Following culture, cellular devices were fixed using 4% paraformaldehyde (ThermoFisher, Waltham, MA) with 0.1% Triton-X (Sigma-Aldrich, St. Louis, MO) at 4°C for 2 h, stained with 1:200 phalloidin (DyLight 554 Phalloidin, Cell Signaling Technologies, Danvers, MA) in PBS, and imaged using a laser scanning confocal microscope (10X, LSM880, Zeiss). Z-stack images were acquired and processed using ImageJ. The distance from the gel interface to each cell that migrated into the acellular region was recorded. Results are presented as an average value ± standard error. Average distances between cell identities and polymerization conditions were compared using a homoscedastic two-sample T-test.

### Generating multiscale vascular networks

#### Chamber fabrication

Molds for the device assembly were fabricated using PDMS and biopsy punches of 4 mm and 7 mm diameters (World Precision Instruments, Sarasota, FL). A large well was made in the PDMS with the 7 mm punch, and a scalpel blade was used to carve two notches into the top edge of the well to act as supports for the large vessel molds. The PDMS well was then mounted onto a glass coverslip via oxygen plasma bonding (Harrick Plasma, Ithaca, NY). The well surfaces were functionalized to facilitate collagen gel bonding using standard methods (24, 28). Briefly, wells were incubated with 2% polyethyleneimine (PEI) for 30 min, then washed three times with deionized water, and subsequently incubated with 0.2% glutaraldehyde (GA) for 1 hr followed by three water washes. A separate PDMS cylinder with guide holes was made with the 4 mm punch and was placed in the center of the well to mold the outer acellular collagen ring.

#### Formation and culture of multiscale vascular constructs

Similar methods were used as described above to create two separate collagen gels that were functionally annealed (57, 58). The 4 mm mold was placed in the center of the well and large channel molds were placed on the notches on the outer PDMS support with ends touching the 4 mm mold. Acellular collagen prepolymer solution was added to the outer ring and allowed to polymerize for two minutes. The 4 mm center mold was removed, and HUVEC-laden collagen prepolymer solution was quickly added to the resulting central void. Following complete polymerization of the concentric structure, the channel molds were gently removed from the outer collagen ring and HUVECs were backfill-seeded at a density of 15 x10^6^ cells/mL into the channels to create cellular large vessels. Devices were cultured at 37°C with 5% CO_2_ for 7 days. Vasculogenesis medium was used to induce plexus formation and was replenished daily until fixation. After culture, multiscale vascular networks were washed three times with PBS, and fixed using 4% paraformaldehyde with 0.1% Triton-X for 2 h at 4°C. Tissues were incubated with anti-CD31 primary antibody (Cell Signaling Technologies, Danvers, MA) overnight at 4°C. Subsequently, samples were incubated with secondary antibody (Donkey anti-Mouse DyLight650, Invitrogen, ThermoFisher) and CF594 conjugated phalloidin (Cell Signaling Technologies, Danvers, MA) overnight at 4°C. Multiscale vascular networks were imaged on a laser scanning confocal microscope (10X, LSM800 and LSM880, Zeiss) using both fluorescence and reflectance imaging modalities. Networks were imaged approximately 100 *μ*m into the gel and large tile scans were acquired to assess the connectivity between the self-assembled network and the large, patterned vessels. Image stacks were auto-segmented using median filtration and thresholding prior to surface generation for 3D visualization with Amira software (ThermoFisher).

#### Vascular network perfusion and flow characterization

Networks were perfused with 410 nm Dragon Green fluorescent microspheres (1% v/v; Bangs Laboratories, Fishers, IN) in culture medium. The microsphere suspension was introduced into one large vessel with a pipette and perfused via a hydrostatic pressure head. Networks were imaged once per second on a laser scanning confocal microscope (5X objective, LSM880, Zeiss) to capture bead movement and cell boundaries. Particle image velocimetry was used to quantify flow using PIV lab plugin (version 2.36.5) on MatLab (R2020b) (59).

## Results

### A multiscale, closed-network vascularized hydrogel

We focused on design requirements to solve two key challenges in creating scaffolds with hierarchically perfused endothelial networks, namely: 1) the interface between self-assembled and deterministically patterned vessels and 2) the “leaky” boundary problem. As such, we micromolded a collagen gel around 3D-printed pins to generate large vessels and used endothelial self-assembly to create a plexus within an acellular boundary collagen gel. Using straightforward and low-cost methods, we were able to rapidly produce multiscale, self-contained vascularized tissue constructs (**Figure 1A**). A PDMS well bonded to a glass coverslip was chemically functionalized with PEI and glutaraldehyde to couple the PDMS walls to the collagen gel, ensuring cell-generated forces would not contract the overall structure of the construct. A PDMS cylinder with two concave guide holes was placed in the center of the well as a mold, and two 3D printed cylindrical, plastic pins (1.5 mm diameter, tapered to a rounded point) were placed in the guide holes to diagonally span between the central PDMS cylinder and the edge of the well within carved notches (**Figure 1B**). The acellular gel was injected into the outer ring created by the mold and allowed to polymerize at 37°C for 2 min (**Figure 1Ci**). The central PDMS cylinder was then removed using forceps, and a HUVEC-laden cellular collagen prepolymer solution was added into the center to polymerize for 20 min at 37°C (**Figure 1Cii-iii**). Following polymerization, vasculogenesis medium was added to the top of the well and the construct was incubated overnight at 37°C (12-18 hrs). To complete endothelialization of the large, deterministic vessels, media was aspirated from the construct and the plastic pins were removed, generating large cylindrical channels in the acellular collagen gel that were in contact the cellular gel (**Figure 1Civ**). HUVECs suspended in culture medium were directly added to these channels and incubated for 20 min at 37°C to allow cells to settle and begin to adhere (**Figure 1Cv-vi**). Following cell seeding, the collagen gels were cultured for 7 days without flow to establish an interconnected, patent, and perfusable vascular construct. Studies conducted herein simply used fluid addition with a pipette and hydrostatic pressure to perfuse the vascular construct; however, if perfusion with a pump is desired, a simple cassette can be created (**Figure 1D**).

**Fig. 1.**
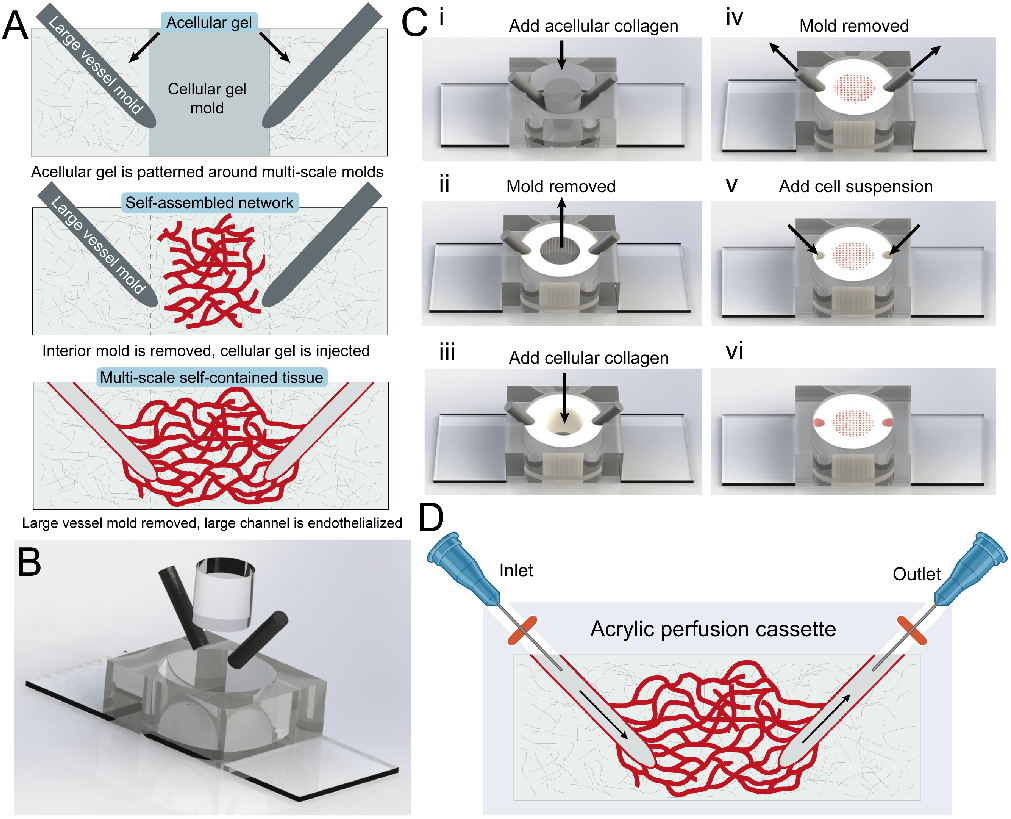
Design and fabrication of a multiscale serially perfused vasculature construct. (A) Cross-section schematic of tissue construct manufacturing set-up. (B) Rendering of a device mold (exploded view). (C) Procedure for creation of tissue construct. i. Inject acellular collagen into outer ring. ii. Remove inner PDMS pillar. iii. Inject cell-laden collagen into the center. iv. Remove large vessel molds. v. Backfill molded channels with endothelial cell suspension. vi. Culture device. (D) Cross-section schematic of interfacing with the large patterned vessels to perfuse the tissue via external pump if desired.

### Creating an annealed and penetrable cellular-acellular collagen gel interface

We solved the “leaky” boundary problem by generating a plexus within an acellular gel. This inhibited the ability of the self-assembled network to form open outlets at the outer boundary of the construct. However, given that the goal of this model is to enable the investigation of vascular network remodeling, we sought to ensure that the cellular-acellular gel interface was fully annealed and allowed cells to migrate across the interface. We hypothesized that a time lag between the polymerization of each gel would affect the formation of the interface between the two gels. To test the gelation times that produced an annealed interface, an acellular collagen prepolymer solution was added to one compartment of a PDMS well and allowed to polymerize for a specified time (0.5, 1, 10, and 30 min). At the end of the time window, a spacing block was removed from the well and another acellular collagen prepolymer solution was added to the resulting open space next to the first collagen gel. In our trials, when the second collagen solution was added 10 or 30 min after the first collagen solution, the interface of the two gels formed an observable seam denoted by accumulated and aligned collagen along the interface as visualized with reflectance confocal microscopy (**Figure 2**). Conversely, when a second collagen prepolymer solution was added 0.5 min after the first gel, no apparent changes in collagen signal nor alignment were observed at the interface compared to the bulk gel. The seam started to become visually apparent by 1 min of delay, revealing that short delay times are needed for fully annealed interfaces to create a monolithic collagen gel, with a stronger collagen alignment and organization along the seam at longer delay times. We used cellular-acellular gel tests to determine if the presence of these varied interfaces would affect local assembly of self-assembled vessels or cell migration into from cellular to acellular gel regions.

**Fig. 2.**
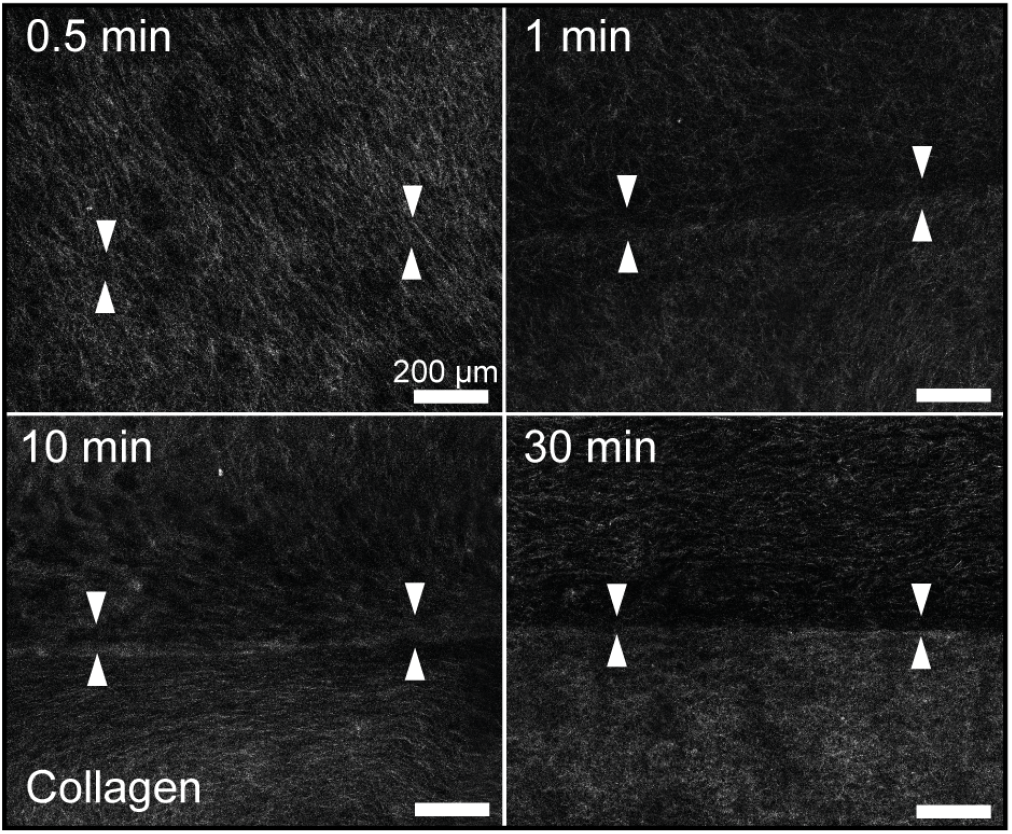
Reflectance confocal microscopy imaging of gel-gel interfaces at four delay time points before adding the second collagen prepolymer solution to the first collagen solution.

When seeded with endothelial cells (**Figure 3A**) stable patterning of the gel regions was accomplished. HUVECs could migrate across the gel interface and vasculogenesis was not effected by the gel-gel interface for all delay times tested. 3T3 fibroblasts were also used as an exemplar of relevant cells with a more migratory phenotype (**Figure 3B**). Co-culture of HUVECs with fibroblasts is known to support plexus formation via paracrine signaling and aid in integration of the vascular plexus with host tissue (36, 60, 61). Similarly, fibroblasts were able to cross the interface into the acellular collage gel at all time delays (**Figure 3C**) and migrate significantly further into the acellular gel than HUVECs (**Figure 3D**). The potential differences in fibroblast migration distance with delay time may be due to a durotactic effect. Neither cell type exhibited cell accumulation or apparent directional migration along the interface for all delay times. These studies confirm that the fabrication methods used will not locally affect cell behavior at gel-gel interfaces and define an acellular region thickness on the order of 0.5-1mm for successful isolation of self-assembled vascular networks.

**Fig. 3.**
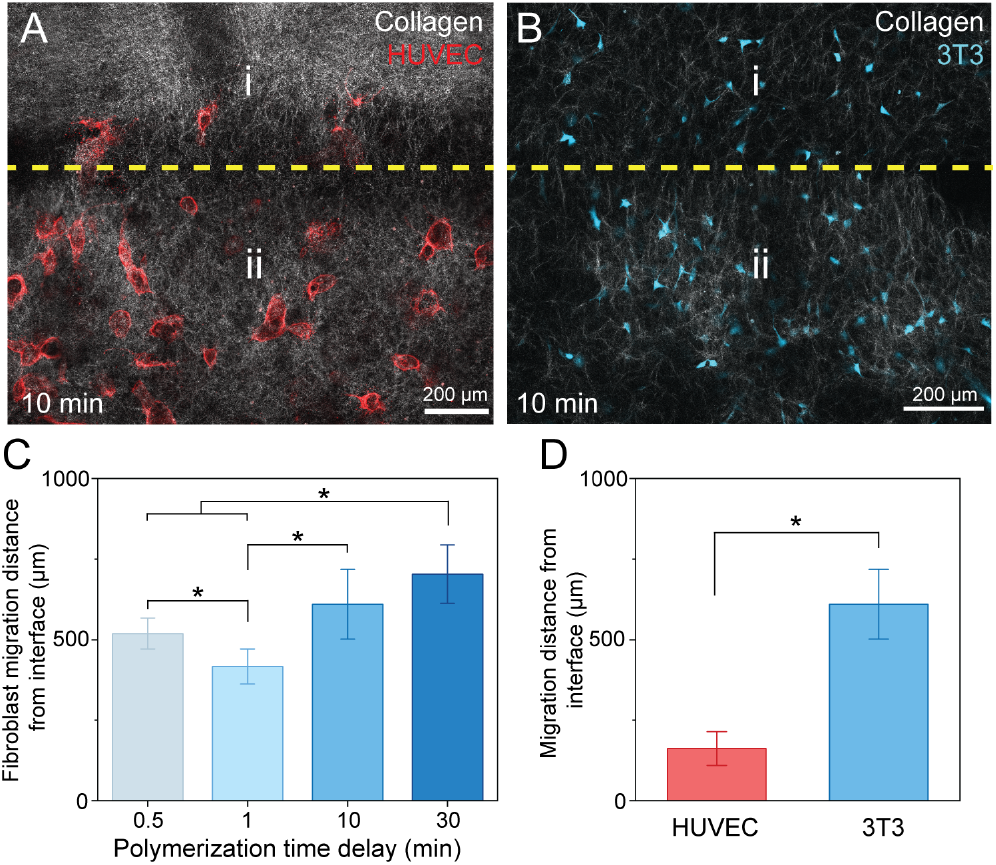
The gel-gel interface allows for cell migration. (A) HUVECs and (B) fibroblasts at the interface (dashed yellow line) following 5 days of culture. Images of the interface between an acellular gel (top, i) and cell-laden gel (bottom, ii) with a polymerization delay of 10 min. (C) Quantification of fibroblast migration into the acellular gel after 5 days of culture for varying delay times. (D) Comparison of HUVEC (n = 10) and 3T3 (n > 25) migration distances into the acellular gel in the 10-minute delay condition. p*<0.05, mean +/- SEM.

### Multiscale connectivity of vascular tissue

We cultured the model without flow, replacing culture medium daily, over seven days to allow for self-assembly of the plexus and formation of a competent endothelial monolayer to line the large deterministically patterned channels. End-point staining with PECAM-1 and phalloidin revealed the maintenance of a cell-free collagen region along the outer boundary of the construct with a robust self-assembled vascular plexus within the central region of the collagen gel consistent with previous studies (28, 35, 46, 62). Confocal z-stack imaging was used to visualize the connectivity between the self-assembled vessels and large deterministically patterned vessels within the constructs. The patterning and connectivity of the multiscale vessels were assessed in the (X, Z) (**Figure 4A**) and the (X, Y) (**Figure 4B**) imaging planes. In all constructs tested, the self-assembled network produced robust vessels that visually anastomosed to the large, endothelialized, patterned vessels at several discreet locations along the channel. Immunostaining confirmed the junctional localization of PECAM-1 in the large patterned vessels indicating proper cell phenotype (**Figure 4C**). To aid in the visualization of the network’s 3D structure and hierarchical connectivity, Amira software was used for vascular segmentation and 3D rendering (**Figure 4D**). At high magnification active sprouting from the large, patterned vessel and integration between the self-assembled network and large vessel was observed (**Figure 4E**).

**Fig. 4.**
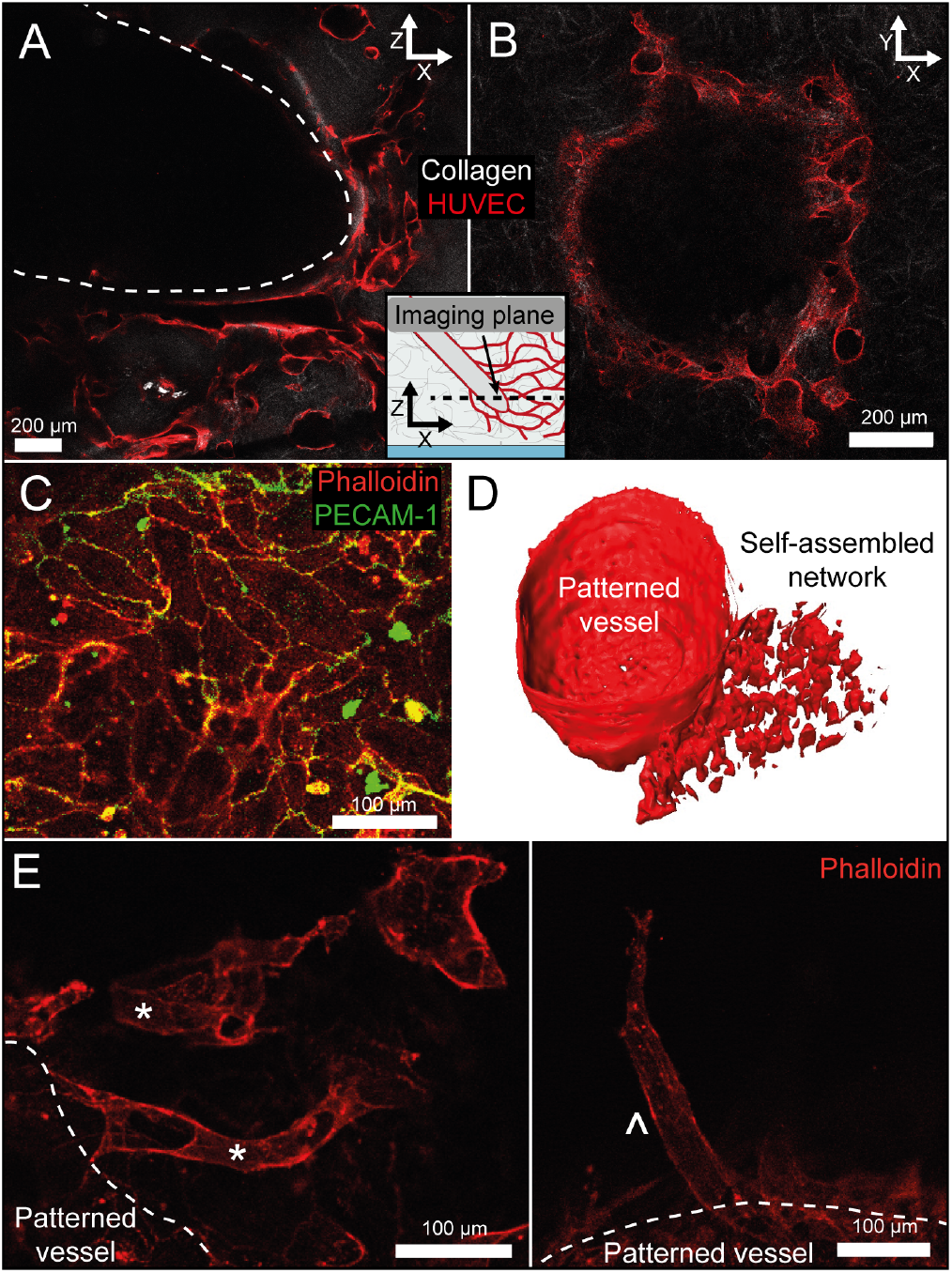
Multiscale connectivity between the large, micropatterned endothelialized channels and the self-assembled network. Large, patterned vessels were integrated into the self-assembled network, as seen in (A) X/Z and (B) X/Y imaging planes. (C) Immunofluorescent staining for PECAM-1 and F-actin in the large vessel confirmed proper cellular phenotype. (D) 3D reconstruction of a region around a large patterned vessel in a multiscale vascular construct. (E) Higher magnification imaging of the constructs showed clear integration between length scales (*\**) and sprouting from the large vessels (∧).

### Perfusion through the network

To determine the connectivity and patency of the multiscale network, 140 nm fluorescent microspheres suspended in cell culture medium were perfused through one of the large patterned vessels into the network using a hydrostatic pressure head. To enable live imaging of the complete vascular network during perfusion, mCherry expressing HUVECs were used for these studies (**Figure 5A**). Using fluorescence confocal microscopy, time series images were taken for approximately one minute after introduction of the pressure head. The microspheres were seen flowing from the large vessel through the self-assembled network (**Figure 5B**) into the outlet solely within the lumen of vessels **(Figure 5C**). To confirm the microspheres flowed unidirectionally through the network, particle image velocimetry (PIV) was performed on an image section with the inlet and self-assembled network in frame (**Figure 5D**). These data validate not only the multiscale connectivity of the construct, but also successful development of a perfusable, self-contained vascular system without “leaky” vessels or boundaries.

**Fig. 5.**
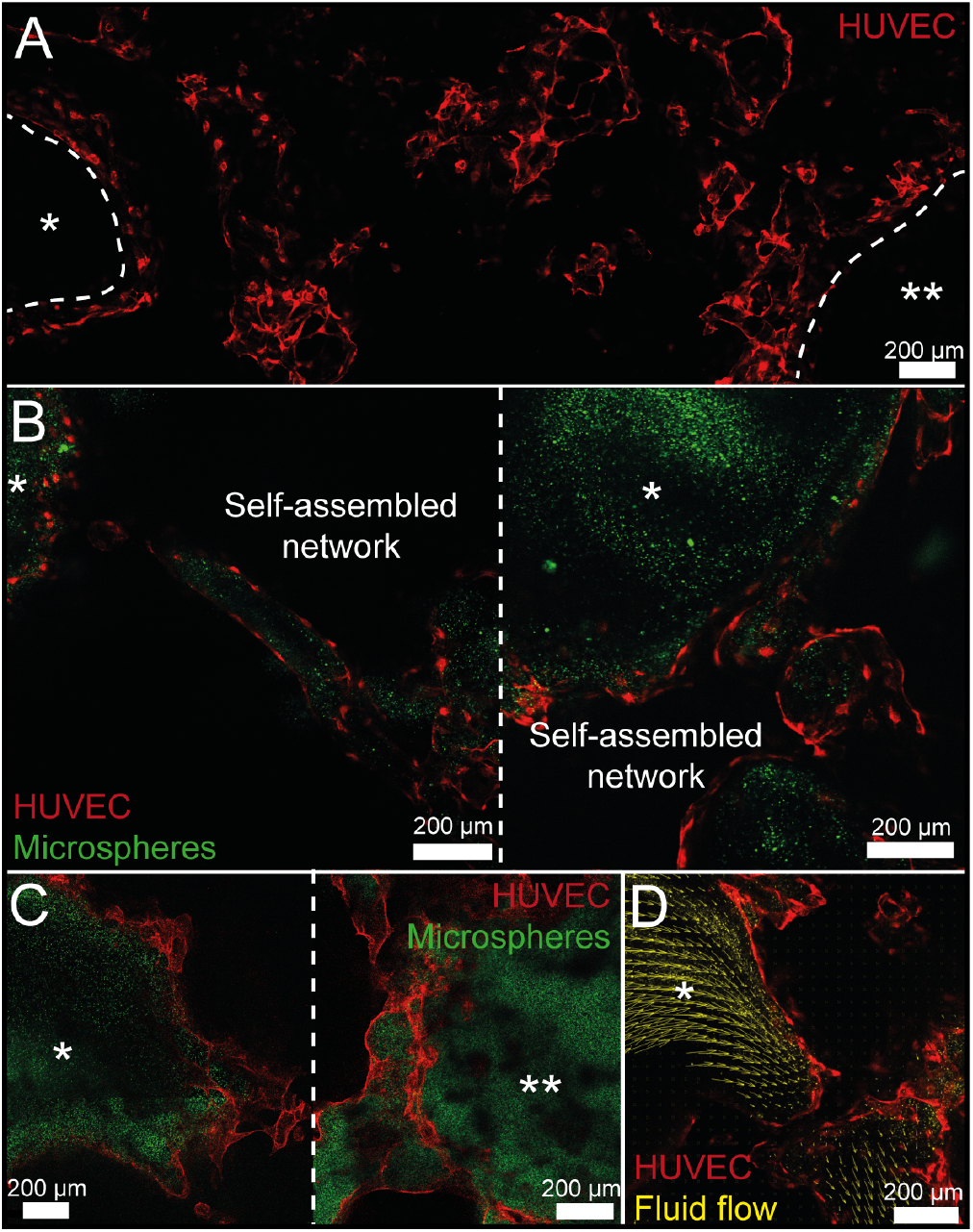
Multiscale construct is perfusable and does not leak at hydrogel boundaries. (A) *In situ* imaging of a single focal plane of a tissue construct with HUVECs expressing mCherry. Timelapse imaging of 410 nm fluorescent microspheres (green) perfusion through construct with fluorescent HUVECs (red) of (B) multiple construct inlets (*) and (C) the inlet (*) and outlet (**) of the same construct demonstrates full connectivity of the multiscale vascular network. (D) PIV of the microspheres in the construct quantifies vascular fluid flows.

## Discussion

The vasculature develops through a complex series of coordinated events to generate a mature, hierarchically organized vascular network (63). The two main mechanisms for the establishment and growth of the vasculature are vasculogenesis, the formation of new vessels from endothelial cell aggregates to generate an initial primitive stochastic vascular network (64), and different modes of angiogenesis, the formation of new vessels from existing vessels (25, 65, 66). The formation of a mature tissue vasculature during development requires extensive use and coordination of the dynamic processes of growth, remodeling, and pruning. The creation and then subsequent refinement and remodeling of a self-assembled vascular plexus results in a highly efficient hierarchical architecture. However, all vascular networks are not the same; studies have demonstrated that microvascular networks are distinct within different tissues and vary with sex (67–70). Remodeling of vascular networks is, in part, regulated by fluid flow and the local mechanical microenvironment and these factors continue to play a significant role in proper endothelial function throughout the entirety of the vessel lifespan (71–73). Understanding the underlying mechanisms and endothelial-stromal cell interactions associated with the remodeling and maintenance of vasculature is a highly active area of biological research. Whereas *in vivo* models are the standard for vascular network investigation, an *in vitro* model system that incorporates a physiologically relevant vessel architecture and control over biomaterial properties, stromal cell identity, spatial organization, and fluid flow would enable a mechanistic understanding of vascular network development, homeostasis, and remodeling. Ultimately, these data would inform optimal vascular network design for engineered tissues and tissue flaps in addition to mechanistic targets for human vascular disease.

Several approaches have been employed to develop a hierarchical vascular network *in vitro*, however, hurdles still exist for achieving model relevance and translational capability. The incorporation of structured (e.g., deterministically engineered, etc.) and self-assembled (e.g., stochastic plexus assembly, etc.) vessels is necessary to produce multiscale, functional vasculature with hierarchical structure into self-contained engineered tissues. Combined, these approaches offer the ability to self-assemble microvascular architectures rapidly, regionally, or across large tissue volumes, coupled with precise spatial control and architecture of large printed or patterned convective vessels. Hybrid approaches, if successfully implemented, offer a path forward for creating multiscale perfusable networks for vascular development, tissue models, and clinical translation.

To this end, we have developed a multiscale vasculature construct that integrates both bottom-up and top-down techniques to create a serially perfusable, hierarchical vascular network. A frequent problem of self-assembly is leaky boundaries, or open lumens at the hydrogel boundary. To overcome this, our system encapsulated the cellular gel region within an acellular gel (**Figure 1**), which constrained the self-assembled microvascular network to the interior of the construct and effectively sealed the boundaries. Studies investigating the acellular-cellular collagen gel interface confirmed an annealed interface between the two gels when the tissue construct was fabricated quickly (**Figure 2**). Longer assembly delay times alter the collagen alignment and accumulation at the gel-gel interface; however, these microstructural changes did not influence the ability of fibroblasts or HUVECs to migrate into the acellular region or alter microvascular formation across the boundary (**Figure 3**). Large-scale self-assembled microvascular networks were formed across the entirety of the 4 mm inner cellular region, and patterned channels were created and anastomosed to the self-assembled network (**Figure 4**). Fluid flow was introduced into the plexus via hydrostatic pressure gradients from pipetted fluid at the large, patterned vessel inlet (**Figure 5**). Scaling of the microvascular network size and geometry is possible with changes to the fabrication mold and blunt-end needles integrated into a simple cassette could be used for large vessel perfusion via external tubing and pumps (**Figure 1D**). Due to the system design the construct could be live imaged *in situ* for quantification of fluid flow (**Figure 5D**). Together, these properties create a simple to fabricate, inexpensive model to study vascular remodeling and maintenance mechanistically.

This simple construct can be optimized and advanced to add further complexity to the vascular network or for use in other applications. These methods are compatible with 3D bioprinting and other advanced methods of creating deterministic large vessels, which can be used to add complexity to the hierarchical vascular network. Supporting stromal cells could be added to the cellular gel when initially creating the plexus to dissect the role of individual stromal cell populations and reciprocal signaling between cell populations in the remodeling and maintenance of vasculature. Additionally, the use of straightforward fabrication methods and the ability to create this system without the need for specialized microfluidic devices expand the applicability and translational potential for this *in vitro* vascular model. Devices and methods that are intentionally designed to be made with common benchtop equipment and inexpensive materials enable broad democratization of these methods, which increases the knowledge and research potential across fields (74). Overall, the methods developed here are widely applicable, easy to incorporate, and can act as a building block for many other self-contained engineered tissues to enhance the complexity and physiological relevance of such models.

## Conclusion

We developed an approach to create a serially perfusable, multiscale vascular network using simple and easily accessible methods. A cellular gel within an acellular gel successfully creates a closed network without impacting vascular self-assembly or cellular migration. This construct results in a vascular network that can be serially perfused without complex microfluidic techniques and imaged *in situ* to quantify fluid flows throughout the network for direct mechanistic investigation of vasculogenesis, angiogenesis, and vascular remodeling.

## ACKNOWLEDGEMENTS

The authors thank Jasmine Shirazi for assistance with collagen interface imaging and Katherine M. Nelson, Ph.D., for reviewing and commenting on the manuscript. This work was supported in part by grants from the University of Delaware Research Foundation and the National Institutes of Health: R01DE029655 and P30GM110758. The BioRxiv template was adapted from the Henriques lab.

## AUTHOR CONTRIBUTIONS

Conceptualization (MJD, JPG), Methodology (MJD, JPG), Investigation (MJD, OM, BC, DCR), Validation (MJD, OM, BC, JPG), Visualization (MJD, OM, BC, JPG), Formal Analysis (MJD, OM, BC, JPG), Writing – Original Draft (MJD), Writing – Review & Editing (MJD, OM, BC, DCR, JPG), Supervision (JPG), Project Administration (JPG), Funding acquisition (JPG)

